# Writing for identity? Exploring the motivations of pre-college students to participate in science publication

**DOI:** 10.1101/2025.03.27.645794

**Authors:** Sarah C. Fankhauser, Tanya Bhagatwala

## Abstract

Scientific publication represents a pinnacle of professional communication in science. While traditional pre-college science classrooms offer various outlets for scientific communication through lab reports and presentations, access to scientific publication, in particular, remains largely out of reach for these students. Yet, this landscape is beginning to shift as an emerging cohort of young researchers challenges this barrier by publishing their work in peer-reviewed journals. This study investigates why pre-college students voluntarily undertake the rigorous process of scientific publication, revealing important insights about student motivation and identity development in science. Through analysis of survey responses from student authors using the lens of science identity theory, we discovered that students view publication as a vehicle for contributing to the scientific community and building one’s own self-efficacy in the process. Female-identifying students in particular emphasized publication as a pathway to gain recognition as scientists and develop professional writing skills. These findings challenge traditional assumptions about pre-college science communication and suggest that access to authentic publication opportunities may be a powerful, yet underutilized tool for fostering science identity and expanding participation in STEM fields. Our research provides critical insights for educators and policymakers seeking to create more authentic and empowering pathways into science careers.

## Introduction

Ten years have passed since the Next Generation Science Standards (NGSS) were released and set the framework of practices meant to prepare students in their 21st century skills and understanding of science (NGSS Lead States, 2013). NGSS emphasizes engaging students in authentic *practices* that help students develop their scientific literacy and transfer skills across disciplines. The eight practices articulated in the Next Generation Science Standards have become key components of the science curriculum for the K-12 classrooms. Among these practices, Communicating Information (Practice 8) stands out as particularly crucial, not just for developing scientific understanding, but for fostering scientific identity and inclusion in the broader science community (Kelp et al., 2024).

As defined by NGSS, students may engage in authentic scientific practices through creating research questions, designing experiments, explaining findings, or reading the literature (NGSS Lead States, 2013; Chapman and Feldman, 2017). And the impact of such authentic engagement can be powerful as demonstrated in a study by Chapman and Feldman (2017), in which urban high school students participated in a 6-week science research project in collaboration with a local university professor. Students who viewed the project as an authentic research program were more likely to report higher science identity. After completing the research project, students also demonstrated a more diverse view of *who* could be a scientist, suggesting that such authentic experiences can broaden one’s perspective on what it means to be a scientist (Chapman and Feldman, 2017).

Such authentic experimental experiences are becoming more common in high school classrooms, however there is less research regarding authentic communication practices at the high school level. In a typical research experience, communication outputs may be operationalized through lab reports, posters, reflective writing, research proposals, etc. (Jacquez et al., 2020; Baker et al., 2009; Vries et al., 2019; Watts-Taffe et al., 2022). However, a growing number of students are engaging in more complex and professional communication endeavors, including scientific publication. Previous evidence with early-career scientists showed a positive relationship between interest in science communication tasks and expected positive career outcomes, and science communication self-efficacy was one variable that significantly predicted career intention (Anderson, et al., 2016). More recently, a longitudinal study revealed that science communication skills significantly contributed to a trainee’s intention to persist in research (Cameron, et al., 2020). While this evidence is for more advanced students, the data suggests that engaging students in science communication can be an influential factor in shaping their interest in, and intention to pursue, science.

Given the demonstrated relationship between science communication and persistence in science, and the integral part that science communication plays within scientific practice, engaging students early in authentic communication experiences represents a critical educational opportunity. Over the the past decade, there has been a notable increase in venues for pre-collegel students to submit their research papers, receive peer-reviewed feedback, and publish on platforms designed specifically for this population of students(Authors 2018, 2022a). The opportunity to write a research paper and participate in the peer-review and publication processes may provide students with the chance to integrate several NGSS practices, while simultaneously fostering their interest in science and strengthening their scientific identities (Authors, 2018; Authors 2021).

### Program Overview: [Journal]

One such journal which provides such publication opportunities to middle and high school students is the [Journal. The [Journal] provides middle and high school students with an authentic introduction to scientific publication through a supportive, scaffolded peer review process. Founded in 2011 as an open-access journal, [journal] has grown from publishing exclusively biological sciences to now accepting hypothesis-driven research across all STEM fields. The journal’s mission centers on making scholarly publications accessible to young scientists by providing the necessary tools, mentorship, and community support.

Students typically conduct their research through school programs, science fairs, or independent projects under the guidance of teachers, parents, or professional scientists. When ready to publish, students prepare manuscripts following standard scientific article formatting, including introduction, methods, results, and discussion sections. The journal provides extensive guides and models through its website to help students navigate this process. Upon submission, manuscripts undergo a structured review process designed to mirror professional academic publishing while maintaining appropriate support for young authors. Papers first go through pre-review to ensure basic components are present. Then, each manuscript is assigned to 3-4 expert reviewers, typically graduate students, who provide detailed feedback on both scientific content and communication. Reviewers, who must complete training and pass a qualification quiz, are instructed to offer constructive, supportive feedback rather than pure criticism. This approach helps students learn from the process rather than feeling discouraged (Otero, 2022).

The revision process forms the heart of the learning experience. An editor synthesizes reviewer feedback into a comprehensive editorial letter, guiding students through needed improvements (Authors, 2016, 2022). Rather than rejecting papers (except in cases of plagiarism or ethical issues), the process focuses on helping students develop their work to meet publication standards. That being said, we do have about 30% of students who submit papers decline to resubmit following the receipt of the reviews. Once the scientific content meets requirements, manuscripts move to copyediting and final proofing. This extensive process creates meaningful opportunities for students to engage with professional scientists through the review process, learning firsthand how scientific knowledge is constructed and communicated. By emphasizing learning through revision and scientific discourse rather than gatekeeping, [journal] makes scholarly publication accessible to young researchers while maintaining the rigor of the scientific process.

This comprehensive approach has yielded impressive results. As of 2025, [journal] has published over 1300 student articles. More importantly, research shows that participating students demonstrate increased confidence in scientific writing and stronger science identity (Authors, 2021; Authors, 2022a). The journal manages this substantial workflow through an online submission portal while maintaining high standards through a network of volunteer graduate students, postdocs, and professionals who serve as reviewers (Authors 2016).

To support broader implementation of writing resources or using the published articles within the classroom, [journal] provides resources to help teachers integrate [Journal] processes and/or papers into their science curriculum (Authors, 2022b). The goal is that professional science communication becomes a standard part of the science curriculum that engages students in the aspects of knowledge-construction that they otherwise would not encounter until college or after. Our innovative approach demonstrates how pre-college students can meaningfully participate in scientific publication when given appropriate support and scaffolding. Through careful attention to both process and mentorship, [journal] helps develop the next generation of scientists by engaging them in authentic scientific communication early in their academic careers.

## Theoretical Framework

Understanding why pre-college students voluntarily engage in scientific publication requires examining both their motivations and how the publication process may shape their developing scientific identities. While scientists spend a considerable amount of time engaging with scientific literature and communication, these practices are notably absent from pre-college science education. Hence, a significant gap exists between classroom science and authentic scientific practice, suggesting the presence of a “hidden curriculum” in science. This makes participation in scientific publication particularly intriguing because, despite the lack of education on disciplinary literacy skills, these students still choose to engage in a challenging process that extends far beyond typical academic requirements.

### Science Identity as a Framework for Understanding Motivation

In this investigation, we draw on the theory of science identity development to examine students’ motivations to participate in the publication of their research (Figure 1). The theory of science identity has been examined by Carlone and Johnson (2007), who used the concept of identity posited by James Gee to develop their model. According to Carlone and Johnson, science identity is influenced by the three interconnected dimensions of “performance”, “competence”, and “recognition.” By articulating these three dimensions, the authors recognize that identity development includes both the ability *to do* the actions of science and to be seen by others as a “science person.”

**Fig. 1.**
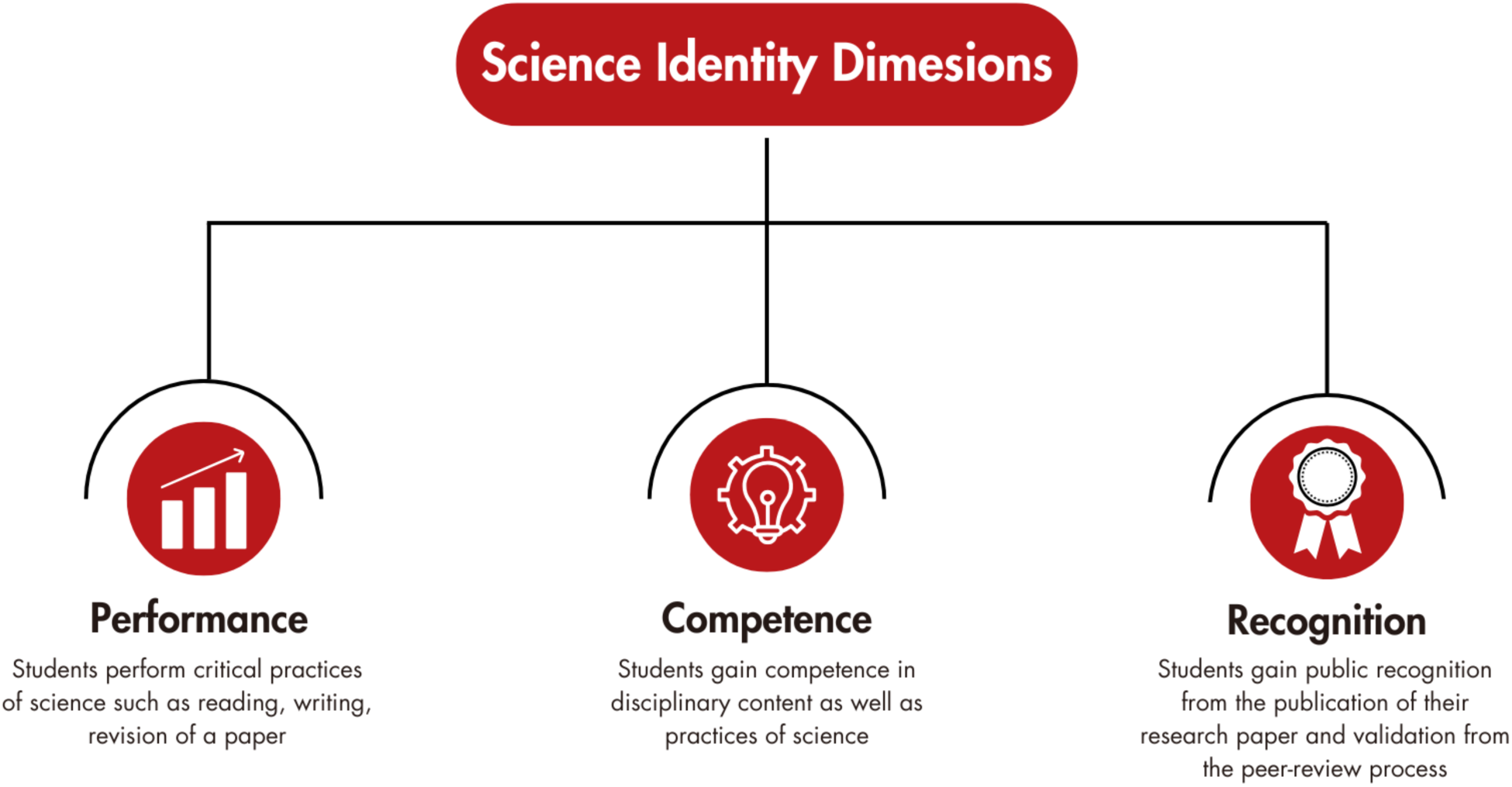
Three dimensions of science identity (above) paired with the outcomes likely achieved when students participate in the publication of their science research papers (below)

In their model, Carlone and Johnson conceptualize identity development as partly situational, that is dependent on the social and logistical structures present to support students in developing their scientific skills. Without opportunities to actively practice science (e.g. through conducting experiments or communicating findings), it would be difficult for students to develop a science identity. Other researchers have used this model to understand how engagement in authentic science practices facilitates students’ science identity and perceptions of who can be a scientist (Chapman and Feldman, 2017; Huffmyer, et al., 2022. What has become clear is that science identity formation is a process, and one’s science identity can be strengthened or destabilized with positive or negative experiences (Robinson et al., 2018 and 2019).

The science identity framework is particularly relevant for understanding student motivation to publish because the act of scientific publication inherently engages all three dimensions:

#### Performance

Publication requires students to demonstrate their ability to engage in authentic scientific practices - not just conducting research, but also communicating that research in ways that align with disciplinary norms. Cameron et al. (2020) have shown that engagement in scientific communication activities serves as a key performance indicator that shapes how early-career researchers see themselves within the scientific community.

#### Competence

Through the publication process, students develop and demonstrate their understanding of both scientific concepts and the processes by which scientific knowledge is constructed and disseminated. Murphy and Kelp (2023) demonstrate that students’ scientific communication skills are intimately connected to their science identity development, suggesting that mastery of scientific writing may be a crucial component of how students assess their own scientific competence.

#### Recognition

Perhaps most significantly, publication provides students with formal recognition from the scientific community through peer review and eventual publication. At the graduate level, a peer-reviewed publication is a key metric for assessing success in academic research, and often provides a catalyst for future research. For pre-college students, peer-review and publication provides external validation to a students’ ideas and research (Authors, 2021) The public nature of publication makes it a powerful vehicle for recognition, as students’ work becomes part of the scientific literature.

#### Operationalizing Science Identity in This Study

This science identity framework is particularly useful for our study given our prior findings that publication influences students’ confidence and self-efficacy in science, which are key elements of “performance” and “competence”. Students also report increased recognition as scientists and belonging within a scientific community following publication of their papers (Authors 2021, 2022, 2023). Notably, interviews with students revealed that students view publication as a mechanism to effect change within their communities (Authors, 2021), thus suggesting that some students may enter the publication process with expectations for recognition and impact—not only by and within the scientific community but also from their broader networks. However, the range and relative salience of students’ initial motivations for pursuing publication remain unexplored. Having a broader understanding of how students perceive and engage with publication through an identity lens may reveal important insights about why some students choose to pursue publication. Furthermore, this framework allows us to examine whether certain dimensions of science identity– such as recognition versus competence– serve as a stronger motivator for specific student populations.

We operationalized the Carlone and Johnson science identity framework in two ways. First, we used the three dimensions to categorize the pre-generated response options in our multiple-choice motivation question. Response options were classified as either “recognition” (e.g., “I want to be recognized as a scientist”) or “competence” (eg, “I want to learn how to write a scientific paper”). Notably, we did not have a response option that could be clearly categorized as “performance”; however, we acknowledge that the boundaries between the categories may not be perfectly distinct, so some options that we categorized as “performance” or “competence” could be viewed as “performance.” Second, we used all three dimensions to deductively code students’ open-ended responses about their motivations. Together, these approaches allow us to identify which aspects of science identity were most salient to students as they entered the publication process.

#### Why Science Identity Over Alternative Frameworks

We considered Eccles’ Expectancy-Value Theory (Eccles & Wigfield, 2002) as an alternative framework, which posits that achievement-related choices are predicted by individuals’ expectations for success (competence beliefs) and the value they place on the task (such as intrinsic and utilitarian). While this framework has been used to understand and predict activity participation in STEM Wang and Degol, 2013), we ultimately selected science identity as the more appropriate framework to analyze our data. First, our past work indicated that students perceived publication as influencing their confidence, self-efficacy, and sense of belonging within science (Authors 2022, 2023), suggesting identity development as central to their experience. Second, Expectancy-Value Theory focuses primarily on predicting future choices, whereas we were interested in understanding motivations for a choice students had already made. Third, science identity theory better captures the social and relational aspects of scientific practice—particularly the dimension of recognition—which our prior interviews suggested was important to students who saw publication as a mechanism to effect change within their communities (Authors, 2021).

### Research Questions

The framework of science identity is particularly useful for our study because publication encompasses each dimension of science identity while also highlighting potential gender differences in how students approach and value different aspects of the publication process. Given that academic writing and publication are generally part of the “hidden curriculum” in science (Moradi et al., 2023; Anlar & Phillips, 2023), understanding how students perceive and engage with publication through an identity lens may reveal important insights about why some students choose to pursue publication while others do not. Moreover, this framework allows us to examine whether certain dimensions of science identity (such as recognition) may be more salient motivators for particular groups of students. Thus our research questions are:

1. To what extent are students motivated to engage in publication for accomplishment and/or impact (recognition), as a way to develop scientific skills (competence), or as part of “dong science” (performance)?
2. Do motivations differ based on the self-reported gender of the student respondent?

## Methods

*Ethics and Consent:* This study was approved by the Emory University Institutional Review Board (IRB STUDY00000797). Students provided informed assent to participate in the research. Parental consent was waived by the IRB as the study posed no more than minimal risk to participants.

### Survey Development and Administration

The survey instrument was designed to assess student authors’ experiences with peer review and publication, drawing on established research frameworks. The survey incorporated validated items from the Persistence in the Sciences (PITS) survey (Hanaueret al., 2016), which includes Likert-scale items assessing project ownership and scientific community values, as well as Likert-scale items measuring self-efficacy and science identity.

The survey was developed with two primary aims: (1) to generate knowledge about the value of peer review and publication on scientific inquiry skills, STEM self-efficacy, and sense of community; and (2) to understand how participation in peer review and publication is impacted by experiential and contextual factors, including the role of research mentors. Some of this work has been published previously (Authors, et al 2022; Authors, et al 2023)

Motivation questions were specifically developed by the research team to explore why students chose to engage in the process of writing and publishing their scientific research. The survey was designed to be administered at two time points (pre- and post-publication) to capture changes in students’ perceptions and beliefs throughout the publication process.

A focus group of student authors convened in 2020 to provide feedback on the survey instrument, particularly on the interpretation and wording of questions. Based on this feedback, the motivation question was modified to use a multi-select format, and the wording of some response options was refined to better capture student perspectives.

The survey was administered through Qualtrics to all student authors who submitted papers to [Journal] between November 2020 and June 2022. Students received the survey link via email immediately following their paper submission. To incentivize participation, respondents could receive a $5 Amazon gift card upon completion of a subsequent Google form. All survey responses were collected anonymously. Of the 613 students who submitted papers during this period, 254 completed at least part of the survey, resulting in a 41.9% response rate. Given the anonymous nature of the survey, we are unable to generate any hypothesis on who did not complete the survey and why.

### Quantitative Analysis

For this study, we focused our analysis on select key questions from a larger survey, related to student motivation for publishing (Supplement 1). One question was a multiple-select question to identify their motivation(s) for submitting a paper for publication, which received 170 responses (81 male-identifying, 89 female-identifying). Response options were categorized according to Carlone and Johnson’s (2007) science identity framework. Options were categorized based on their primary alignment with the framework’s dimensions: Recognition items focused on being seen by or connecting with others in the scientific community, as well as gaining external validation. Performance items centered on enacting core scientific practices, particularly the practice of disseminating research findings. Competence items emphasized learning and skill development within scientific domains. Some items could potentially align with multiple dimensions but were categorized based on their primary emphasis as indicated by the action verbs and intended outcomes in each statement (e.g., “want to learn” indicating competence, “want to connect” indicating recognition).

We also asked a pair of multiple-select questions regarding who helped students perform their research and write their manuscripts (with 207 and 208 responses to these questions, respectively). We also asked where students performed the majority of their research ( 208 responses).

Students were asked demographic questions regarding their racial identity, gender identity, and grade-level (Authors, 2022). For this study we focused on gender differences since this gave us a roughly equal number of participants to analyze. The question was phrased as: “What gender do you most identify with?” To determine if there were significant differences in factors between male-identifying and female-identifying students, we conducted two-tailed Z-tests of proportions for each response option. Statistical significance was set at p < 0.05. Non-binary and self-subscribe responses were not included in this comparative analysis due to small sample sizes. We used an alpha level of p < .05 without correction for multiple comparisons.

This decision reflects the exploratory nature of these analyses and balances the risks of Type I versus Type II errors in detecting potential gender differences in an understudied area. To evaluate the practical significance of gender differences, we calculated Cohen’s h as our effect size measure. This statistical approach transforms proportional data to better handle values that are bounded between 0 and 1, making it particularly suitable for comparing the percentage of responses between independent groups (Cohen, 1988; Fritz et al., 2012). Following Cohen’s guidelines, we interpreted effect sizes as small (h=0.20), medium (h=0.50), and large (h=0.80).

### Qualitative Analysis

The second question was an open-response item asking students to elaborate on their selected motivations. We received 89 responses to this question (50 responses from female-identifying students, and 39 from male-identifying students. These responses were analyzed using a multi-phase coding approach:

1. Initial Reading: The primary researcher conducted an initial open-reading of all responses. Upon sharing some initial themes that emerged with other researchers, including at a round table presentation at [conference], she was advised to take a deductive approach, looking for codes that aligned with Expectancy Value (Eccles & Wigfield, 2002) and science identity (Carlone and Johnson, 2007). The Eccles Expectancy-Value is a framework that uses two key factors to determine why individuals engage and persist with a certain task: expectancy for success (belief that the individual will accomplish the task) and task value (the importance or worth of the task). Task value is further broken down into attainment value (importance for one’s identity), intrinsic value (personal interest), utility value (usefulness for goals), and cost (sacrifices required). These are all influenced by cultural, social, and experiential factors and together play a role in students’ choices, motivations, and performance in academic spaces (Eccles & Wigfield, 2002).
2. Initial Deductive Coding: Two initial independent sets of codes were developed based on the theoretical frameworks of expectancy value and science identity. Through iterative reading and coding, the primary author determined that the open response data fit the codes of science identity more so than expectancy value. In particular, it was challenging to disentangle how student responses may fit into the expectancy value code of “utility” versus “attainment”, but the majority of those responses were coded into “recognition” based on science identity. For example, the student response “I wanted validation on my growth as a scientist, to introduce this great piece of medical equipment, and to share my findings with the scientific and medical community” was initially coded as “attainment” using expectancy value theory and “recognition” using science identity theory.
3. Deductive Coding: Following the theoretical framework of science identity, the primary researcher developed a codebook with three primary constructs:

- Recognition: Responses indicating desire to make an impact or be seen as a scientist
- Competence: Responses expressing a desire to improve skills or learn
- Performance: Responses indicating engagement in activities that scientists typically do

The codebook was finalized with clear definitions and example responses for each code (see Table 2 in Results). All open-ended responses were imported into Dedoose qualitative analysis software and coded using this framework. We coded at the level of meaning units, distinct ideas or themes within responses, rather than coding entire responses as single units (Saldaña, 2015). This approach yielded 104 coded segments, as some student responses contained multiple motivations that aligned with different dimensions of science identity. For example, a student might express both a desire for recognition (’I want to be seen as a scientist’) and competence development (’I want to improve my writing skills’) within a single response, resulting in two coded segments from one response.

To ensure coding reliability, we employed the following measures:

- Clear operational definitions were established for each code
- Example responses were included in the codebook for reference
- Code applications were reviewed and discussed among the research team
- Codes were applied consistently across all responses

To determine interrater reliability, a second researcher who had not coded the data originally analyzed a sample of the responses (24/104 responses). Cohen’s Kappa was performed to evaluate the consistency of coding between the researchers. On average, there was 90% agreement across codes and a Cohen Kappa of 0.79, indicating substantial agreement across the coders. The code with the largest area of disagreement was “performance,” in which the original coder had double-coded several statements as “performance” and either “competence” or “recognition”.

## Results

Analysis of our pre-publication survey reveals how pre-college students’ motivations for voluntary scientific publication align with Carlone and Johnson’s science identity framework (Table 1). Students generally selected multiple motivating factors (male-identifying students averaging 5.8 factors; female-identifying students averaging 6.2 factors with no significant difference), suggesting that they view publication as serving multiple roles in their scientific development. Through analysis of both multiple-select responses (n=170) and open-ended explanations (n=89), we examined how students’ motivations aligned with the three dimensions of science identity: recognition, performance, and competence.

**Table 1.**
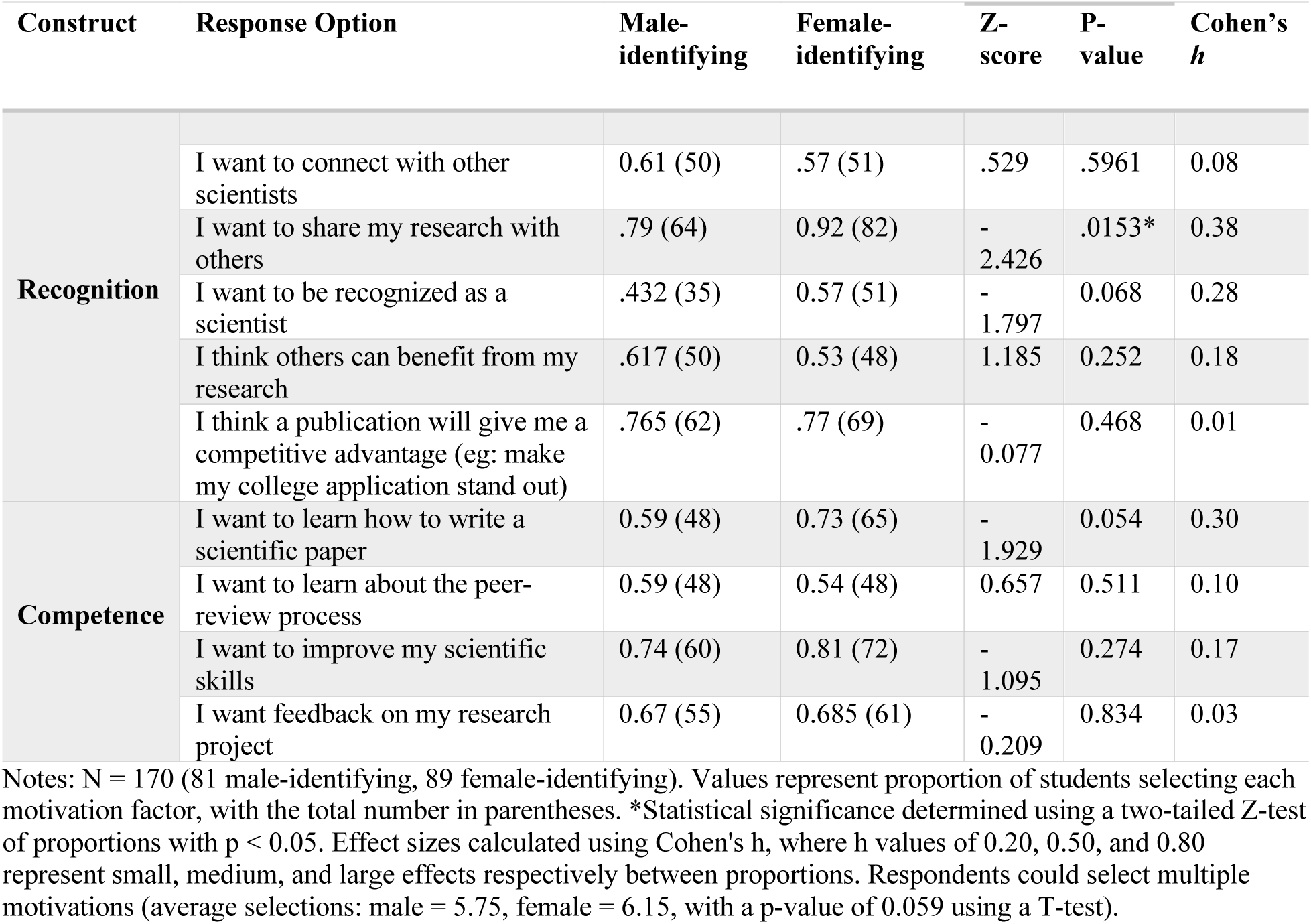
Student Motivations for Scientific Publication by Gender Identity: Response Frequencies and Statistical Comparisons.

### Recognition: Seeking Legitimacy in Science

Recognition emerged as a prominent motivating factor across both quantitative and qualitative data. In the multiple-select responses, the most highly selected option was “I want to share my research with others”, with 92% of female-identifying students and 79% of male-identifying students selecting this option. The gender difference was both statistically significant (p<0.05) and practically meaningful, with the effect size (h=0.38) indicating a small-to-medium difference in how recognition factors into the motivation to publish (Table 1). Female-identifying students were more likely than male-identifying students to select “I want to be recognized as a scientist” as a motivation (57% vs 43%). Though this difference approached but did not reach statistical significance, and the effect size was small (Cohen’s h= 0.28), it suggests a meaningful trend worthy of further investigation.

This emphasis on recognition was further elaborated in open-ended responses, where it emerged as the most prevalent theme (55 coded segments). Students explicitly connected publication to gaining legitimacy within the scientific community. As one student expressed: “Conducting a research project is difficult, but it is very important for me that the process… is veritably credible and respected by the scientific community.” Another student highlighted the transitional aspect of recognition: “I believe that being successful in publishing a science research paper can be a metric to prove my capabilities and attention to detail…I want to show myself that completing such a difficult task is within my reach.”

### Competence: Building Scientific Skills

Students also expressed strong interest in developing their scientific competencies through the publication process. The competence-related motivations focused on skill development, learning about the peer-review process, and receiving feedback on their research. The most frequently selected competence item was “I want to improve my scientific skills”, chosen by 81% of female-identifying students and 74% of male-identifying students. This was followed closely by “I want feedback on my research project”, with 69% of female-identifying students and 67% of male-identifying students selecting this option. While female-identifying students selected competence-related motivations at slightly higher rates for three of the four items, none showed statistically significant differences.

Furthermore, in their open-ended responses (40 coded segments), students elaborated on how they viewed publication as a pathway to developing scientific competence (Table 2). As one student explained: “I want to publish in [Journal] so that I can improve my critical thinking skills… All of which I think will help shape me into a better scientist.” Another student highlighted the learning potential of the review process: “I was drawn to [Journal] specifically because it would be peer-reviewed by senior scientists who are knowledgeable in the field and would provide helpful feedback so I can better my research”

**Table 2.** Coding Framework for Student Publication Motivations Using Science Identity Theory.

| Construct | Description | Example Student Quote |
| --- | --- | --- |
| **Recognition**<br>**(55 segments)** | I want to make an impact/be seen as a scientist- recognition from self or public | "I found [Journal] and wanted to...and be recognized for my work and as an aspiring scientist." |
|  | This will give me an edge for college-recognition by institution | "I would like this competitive advantage to get into the best secondary institute possible so that I can contribute to the future of the world and people" |
| **Competence**<br>**(40 segments)** | I want to improve my skills/learning | "So I wanted to learn about the review process so I can start publishing some work I did in Math and Physics." |
| **Performance**<br>**(9 segments)** | This is what scientists do so I'm going to do it | "Publications serve as milestones in the scientific journey, otherwise, the process of scientific research would seem like an infinite path with no reward." |

### Performance: Engaging in Scientific Practice

The performance dimension of science identity was primarily represented in our survey by the act of constructing or disseminating one’s research. While the multiple-select response, “wanting to share research with others”, could be considered an act of “performance”, we chose to qualify it as “recognition” given that sharing implies an audience, visibility, and community contribution. In the open-ended responses, there were far fewer student responses that aligned with performance (9 coded-segments). However, these comments provided valuable insight into how students view publication as authentic scientific practice. One student noted: “It was the next logical step to take. I went through the program over the summer, performed the experiment, and wrote my sections of the paper.” Such responses suggest these students view publication not just as an add-on activity, but as an integral part of doing science.

### Integration of Identity Dimensions

Many students’ responses demonstrated how these three dimensions of science identity interweave in their motivations for publication. For example, one student wrote: “Publishing in [Journal] would allow be to play a more active role in sharing knowledge within the scientific community and improve my own rigor as an aspiring scientist” This response integrates performance (the act of knowledge dissemination), competence (improving rigor), and recognition (identifying oneself as a scientist).

Notably, even apparently practical motivations like college applications were often framed within the context of scientific identity development. For example, one student noted “I also thought that publishing with [Journal] will give me an advantage for college because it’s something that shows my strong interest in science.”

The pattern of effect sizes observed, with the largest for the recognition item of “I want to share my research with others” (*h*=0.38) as well as “I want to learn how to write a scientific paper” (*h*=0.3) suggests that female-identifying students may place a greater emphasis on formal recognition and skill development through publication.

### Students Self-report Independence in Science

To further understand the context in which students pursued publication, we examined the sources of support students accessed during their research and writing processes. These data provide insight into how students navigate the “hidden curriculum” of scientific publication and may illuminate factors that facilitate or constrain their ability to engage in this voluntary activity.

Students reported a high level of independence in both the research and writing processes. When asked “Who helped you perform the research that you wrote about?”, the most frequently selected response was “Mostly performed independently,” chosen by 38% of both male-identifying (n=38) and female-identifying (n=41) students (Table 3). This pattern of independence persisted into the writing phase, with 44% of male-identifying students (n=44) and 43% of female-identifying students (n=46) reporting that they “mostly performed independently” when writing their papers (Table 4). No significant gender differences emerged in reported independence for either research (z = 0, p = 1.00) or writing (z = 0.145, p = .88).

**Table 3:** Who helped you perform the research - Gender Comparisons with Effect Sizes.

| Support Source | Male-identifying | Female-identifying | Z-score | P-value | Cohen's $h$ |
| --- | --- | --- | --- | --- | --- |
| Science teacher | .25 (25) | .27 (29) | -0.328 | 0.743 | 0.05 |
| Non-science teacher | .07 (7) | .14 (15) | -1.63 | 0.102 | 0.23 |
| College/university professor | .23 (23) | .24 (26) | -0.170 | 0.865 | 0.02 |
| **Parent** | .29 (29) | .21 (23) | 1.334 | 0.182 | 0.19 |
| **Friend or classmate** | .13 (13) | .17 (19) | -0.804 | 0.421 | 0.11 |
| **Mostly performed independently** | .38 (38) | .38 (41) | 0 | 1.00 | 0 |
| **Other** | .15 (15) | .24 (26) | -1.629 | 0.103 | 0.23 |

Notes: N = 207 (99 male-identifying, 108 female-identifying). Values represent the proportion of students selecting each factor, with the total number in parentheses. \*Statistical significance determined using a two-tailed Z-test of proportions with $*p < 0.05$ . Effect sizes calculated using Cohen's $h$ , where $h$ values of 0.20, 0.50, and 0.80 represent small, medium, and large effects, respectively, between proportions. Respondents could select multiple support options.

**Table 4:** Who helped you write the paper (select all that apply) - Gender Comparisons Effect Sizes.

| **Support Source** | **Male-identifying** | **Female-identifying** | **Z-score** | **P-value** | **Cohen's $h$** |
| --- | --- | --- | --- | --- | --- |
| **Science teacher** | 0.19 (19) | 0.22 (24) | -0.533 | 0.593 | 0.07 |
| **Non-science teacher** | 0.07 (7) | 0.13 (14) | 01.429 | 0.153 | 0.20 |
| **College/university professor** | 0.18 (18) | 0.17 (18) | 0.189 | 0.850 | 0.06 |
| **Parent** | 0.18 (18) | 0.20 (22) | -0.366 | 0.714 | 0.05 |
| **Friend or classmate** | 0.18 (18) | 0.27 (29) | -1.544 | 0.123 | 0.22 |
| **Mostly performed independently** | 0.44 (44) | 0.43 (46) | 0.145 | 0.88 | 0.02 |
| **Other** | 0.12 (12) | 0.19 (20) | -1.385 | 0.166 | 0.19 |

Notes: N = 208 (99 male-identifying, 109 female-identifying). Values represent the proportion of students selecting each factor, with the total number in parentheses. \*Statistical significance determined using the Z-test of proportions with $*p < 0.05$ . Effect sizes calculated using Cohen's $h$ , where $h$ values of 0.20, 0.50, and 0.80 represent small, medium, and large effects, respectively, between proportions. Respondents could select multiple support options.

Beyond independence, students drew on diverse support networks, with no single source dominating. For research support, the most frequently selected option, after independent work, were science teachers (27% of female-identifying students, 25% of male-identifying students), parents (21% of female-identifying students, 29% of male-identifying students), and college/university professors (24% of female-identifying students, 23% of male-identifying students). For writing support, after independent work, the distribution was more even across sources, with science teachers (22% of female-identifying students, 19% of male-identifying students), parents (20% of female-identifying students, 18% of male-ideentifying students), college/university professors (17% of female-identifying students; 18% of male-identifying students), and friends or classmates (27% of female-identifying students, 18% of male-identifying students) all selected at similar rates.

Notably, no statistically significant gender differences emerged in any support source for either research or writing (all p > 0.05). While effect sizes were generally negligible to small (Cohen’s h < 0.20), a few patterns approached small effect sizes: females were slightly more likely to report “other” support for both research (h = 0.23) and writing (h = 0.19), and non-science teachers for research (h = 0.23). Students who selected “other” primarily described receiving help from scientific mentors such as graduate students, postdoctoral researchers, or professional scientists, suggesting that some students had access to formal or informal mentorship beyond traditional educational settings.

The majority of students conducted their research at home (59% of female-identifying students, 67% of male-identifying students), with smaller proportions working in their schools (17% of female-identifying students,19% of male-identifying students)) or university/college laboratories (8% of female-identifying students, 6% of male-identifying students), (Table 5). No significant self-reported gender differences emerged in research location. The prevalence of home-based research, combined with students’ reports of working independently and/or seeking support from parents, peers, and external mentors, suggests that many students were conducting independent or small-scale projects outside traditional institutional settings. This context may help explain both the high rates of reported independence and the reliance on informal support networks, as students navigating the publication process may have had limited access to institutional resources or formal guidance on scientific writing.

**Table 5:** Where students performed the majority of their research - Gender Comparisons with Effect Sizes.

| Location | Male-identifying | Female-identifying | Z-score | P-value | Cohen's $h$ |
| --- | --- | --- | --- | --- | --- |
| **At home** | 0.67 (66) | 0.59 (64) | 1.19 | 0.233 | 0.17 |
| **In the school that I attend** | 0.19 (19) | 0.17 (19) | 0.375 | 0.707 | 0.05 |
| **University/college laboratory** | 0.06 (6) | 0.08 (9) | -0.563 | 0.575 | 0.08 |
| **Other** | 0.08 (8) | 0.16 (17) | -1.76 | 0.078 | 0.25 |

Notes: N = 209 (99 male-identifying, 109 female-identifying). Values represent the proportion of students selecting each factor, with the total number in parentheses. \*Statistical significance determined using the Z-test of proportions with $*p < 0.05$ . Effect sizes calculated using Cohen's $h$ , where $h$ values of 0.20, 0.50, and 0.80 represent small, medium, and large effects, respectively, between proportions. Respondents could select multiple support options.

## Discussion

Our study reveals how pre-college students view scientific publication as a pathway for developing their science identity, particularly through gaining recognition and building s. For students already interested in or engaged with science, publication opportunities may serve as a critical mechanism to simultaneously gain recognition, build competence, and demonstrate performance in science – three key factors that Carlone and Johnson (2007) identify as essential for science identity development. Providing students who are motivated to pursue science beyond the classroom with authentic avenues can be a promising strategy for deepening sustainment in STEM participation.

### Recognition in Science Identity Development

The prominence of recognition-oriented motivations in our findings, particularly among female-identifying students, aligns with previous research highlighting recognition as a critical factor in science identity development and persistence (Robnett et al., 2015). Female-identifying students were more likely to pursue publication to share their research with others and be recognized as a scientist, suggesting they may view publication as a formal pathway to legitimate participation in science. This finding is particularly important given that women remain underrepresented in many scientific fields and often face greater barriers to recognition of their scientific contributions (Kim, et al, 2018).

#### Gender Differences in Science Identity Construction Through Publication

The prominence of recognition as a motivating factor should also be understood within the broader social context in which ‘scientist’ is often coded as male (Finson, 2010), and where even high-achieving girls are less likely to see themselves as ‘science kind of people’ compared to boys with similar achievement (Gauthier et al., 2017). Female students’ selection of recognition-related motivations may not simply reflect different individual approaches, but rather their strategic response to systemic constraints that make their scientific capabilities less readily assumed or acknowledged. This is also supported by the greater emphasis female-identifying students place on the motivation factor of “learning to write a scientific paper”. Together, these selections may reflect not merely a desire for skill development, but an awareness that mastery of formal scientific practices—and external validation of that mastery—may be more necessary to establish legitimacy in a field where their scientific identity is less automatically assumed.In this context, publication becomes a formal mechanism to assert and have validated a claim on scientific identity that may be more automatically afforded to male peers.

The observed gender differences in motivation patterns reveal important insights about how female-identifying students may strategically use publication to build their science identities. Female-identifying students’ greater emphasis on both sharing their research (recognition) and learning how to write a scientific paper (competence), while small in effect, suggests that they may view publication as a formal pathway to overcome traditional barriers in science (Ross, et al., 2022). Their higher likelihood of viewing research sharing as a motivating factor suggests they particularly value the opportunity to actively contribute to scientific knowledge through writing and publication.

The gender differences in the competence-related motivation of learning how to write a scientific paper, may reflect female-identifying students’ strategic approach to science identity development. By emphasizing skill development in scientific writing, female-identifying students may be recognizing writing as a crucial scientific practice that can help establish their legitimacy in the field. This attention to developing competence in scientific communication aligns with research showing that women often need more explicit recognition and validation of their scientific capabilities to develop strong science identities (Carlone & Johnson, 2007). In this context, the publication process may serve as a marker of achievement that reinforces belonging in a space where women have historically been, and continue to be, underrecognized (Ross et al., 2022)). This validation may be especially meaningful for sustaining participation and confidence in science fields.

Other researchers have investigated the intersection of gender-identity, identity in science, and persistence in science. While we focus on one aspect of one’s identity, we acknowledge that identity is not created in isolation but shaped by social exposures, experiences, and appraisals of family and community (Hill et al., 2017). Institutional structures, like schools, are important avenues for appraisal, whether positive or negative, and academic role models that can contribute to one’s identity (Hill et al., 2017; Buday et al., 2012). Buday et al found that people with strong scientific achievement at the high school level and who had positive social support were associated with those individuals seeing themselves in scientific careers and actually having a career in a science-related field. Our student participants, both highly independent and achieving, may be perceiving publication as a validating appraisal that could further contribute to their development of seeing themselves as scientists.

#### Integration of Identity Dimensions Through Scientific Publication

Scientific writing and publication are not merely auxiliary skills but rather constitute core scientific practices essential to knowledge construction and dissemination in science. As emphasized in the Next Generation Science Standards (NGSS Lead States, 2013), communicating scientific information is a fundamental practice that helps students develop scientific literacy and transfer skills across disciplines. Our findings suggest that students recognize this, viewing publication as an opportunity to simultaneously enact scientific practices, demonstrate competence, and gain recognition. *How* students gain this understanding, e.g. through experiences or reinforced by mentors, is not fully understood and would make for a very valuable future study.

Although outside the scope of this study, it’s likely that the full publication process itself reinforces the integration of identity dimensions through peer review, where students receive feedback (recognition of their work) that helps them develop their self-efficacy and competence while simultaneously engaging in the professional practices of the scientific community (performance). This creates a cycle where the performance of scientific practices leads to increased competence, which in turn strengthens their recognition as legitimate participants in science. This cyclic reinforcement may be particularly important for building robust science identities that can persist through challenging academic and career trajectories.

Also, while it is possible that students who chose to pursue the process likely possessed relatively higher science identities, our findings suggest that the process of engaging in authentic scientific writing and peer review further deepened their sense of competence and recognition within the science community. Thus, we suggest that publication may function both as an outcome of a student’s scientific identity and as a mechanism to further science identity development.

The formation of scientific identity involves individuals increasingly seeing themselves as “science kind of people”, but it is also influenced by social factors. Based on the social identity theory, identity develops through interactions and recognition one receives from others within the community they strive to be part of (Hogg et al., 1995). For students, being perceived by mentors, peers, and the broader scientific community as capable contributors to the scientific discourse reinforces their sense of belonging in science (Hill et al., 2018). In this way, engaging in authentic peer review and publication practices not only builds competence but also provides social validation that strengthens students’ science identity.

### Limitations

First, our sample represents a self-selecting group of highly motivated students who voluntarily pursued publication and thus likely possess higher levels of science self-efficacy and interest compared to typical pre-college students. This suggests our findings are not generalizable to broader student populations but could have implications for other endeavors, such as science fairs or other competitions, focused on supporting highly motivated student populations. Additionally, in the future we would like to compare our results to students who decline to participate in the publication process. What parts of the process are perceived as the most substantial barriers? Answering these questions will help us continue to break down barriers to sharing scientific knowledge.

Second, our reliance on self-reported data through surveys captures students’ stated motivations but may not fully reveal underlying factors influencing their decisions to publish. While our mixed-methods approach helps mitigate this limitation, observational studies or longitudinal tracking could provide additional insights into how motivations manifest in actual behaviors and outcomes.

Third, our gender analysis was limited to binary gender categories due to small sample sizes for non-binary and self-identifying students (n < 10). This constraint prevents us from understanding the potentially unique experiences and motivations of gender-diverse students in scientific publication. Additionally, the intersection of gender with other identity factors (e.g., race, socioeconomic status) could not be meaningfully analyzed due to sample size limitations. Finally, our study’s focus on a single journal ([Journal]) may not capture variation in student experiences across different publication venues or formats. The journal’s specific support structures and review processes could influence student motivations in ways that might differ in other publication contexts.

Despite these limitations, our findings provide valuable insights into how pre-college students approach scientific publication and its role in science identity development, particularly regarding gender differences in motivation patterns.

### Implications for Science Education and Scientific Community Engagement

Our findings suggest the need to fundamentally reimagine how we integrate scientific communication opportunities into pre-college education. Rather than treating student research as purely educational exercises, educators and those involved in science outreach could create authentic connections between students and the scientific community. This could manifest through student research symposia, school-university partnerships, and mentorship programs that connect students with working scientists. Such opportunities provide scaffolded experiences that help students develop both confidence and competence in scientific communication while gaining recognition from the broader scientific community.

The development of systematic support structures is crucial for making these opportunities sustainable and equitable. This includes professional development for educators in guiding student research and scientific writing. It also involves partnerships between K-12 schools and scientific institutions, as well as the recognition of communication and publication within science education standards.

Creating more opportunities for authentic scientific communication could have broader impacts on diversity in science by providing formal pathways for recognition and community entrance. Moreover, it could enhance scientific literacy through a deeper understanding of how scientific knowledge is constructed and communicated. This approach aligns with NGSS Science Practice 8, Communicating Information, by providing students with concrete opportunities to develop their scientific identities through meaningful communication practices.

### Future Directions

While our study focused on highly motivated students who voluntarily pursued publication, future research should examine how publication opportunities might support science identity development in broader student populations. Additionally, longitudinal studies could investigate how early publication experiences influence long-term persistence in science, particularly for underrepresented groups. Understanding these relationships could help design more effective interventions to support diverse students’ science identity development and persistence in scientific fields.

## Acknowledgements

This material is based upon work supported by the NSF under Grant No. 2010333

## References

Authors 2016

Authors 2021

Authors 2022a

Authors 2022b

Anderson, C. B., Lee, H. Y., Byars-Winston, A., Baldwin, C. D., Cameron, C., & Chang, S. (2015). Assessment of Scientific Communication Self-efficacy, Interest, and Outcome Expectations for Career Development in Academic Medicine. Journal of Career Assessment, 24(1), 182–196. 10.1177/1069072714565780

Anlar, B., & Phillips, H. (2023). Addressing the “hidden curriculum” in political science publishing. Politics & Gender, 19(2), 611–615. 10.1017/S1743923X23000156

Baker, D., Lewis, E., Purzer, S., Watts, N. B., Perkins, G., Uysal, S., Wong, S., Beard, R., & Lang, M. (2009). The Communication in Science Inquiry Project (CISIP): A project to enhance scientific literacy through the creation of science classroom discourse communities. International Journal of Science Education, 31(5), 619–641.

Buday, S. K., Stake, J. E., & Peterson, Z. D. (2012). Gender and the choice of a science career: The impact of social support and possible selves. Sex roles, 66(*3*), 197–209. 10.1007/s11199-011-0015-4

Cameron, C., Lee, H. Y., Anderson, C. B., Trachtenberg, J., & Chang, S. (2020). The role of scientific communication in predicting science identity and research career intention. PLOS ONE, 15(2), Article e0228197. 10.1371/journal.pone.0228197

Carlone, H. B., & Johnson, A. (2007). Understanding the science experiences of successful women of color: Science identity as an analytic lens. Journal of Research in Science Teaching, 44(8), 1187–1218. 10.1002/tea.20237

Chapman, A., Feldman, A. (2017). Cultivation of science identity through authentic science in an urban high school classroom. Cult Stud of Sci Educ, 12, 469–491 10.1007/s11422-015-9723-3

Cohen, J. (1988). Statistical Power Analysis for the Behavioral Sciences (2nd ed.). Routledge. 10.4324/9780203771587

Eccles, J. S., & Wigfield, A. (2002). Motivational beliefs, values, and goals. Annual Review of Psychology, 53(1), 109–132. 10.1146/annurev.psych.53.100901.135153

Finson, K. D. (2002). Drawing a scientist: What we do and do not know after fifty years of drawings. School Science and Mathematics, 102(7), 335–345. 10.1111/j.1949-8594.2002.tb18217.x

Fritz, C. O., Morris, P. E., & Richler, J. J. (2012). Effect size estimates: Current use, calculations, and interpretation. Journal of Experimental Psychology: General, 141(1), 2–18. 10.1037/a0024338

Gauthier, G. R., Hill, P. W., McQuillan, J., Spiegel, A. N., & Diamond, J. (2017). The potential scientist’s dilemma: How the masculinization of science shapes friendships and science job preferences. Social Sciences, 6(1), 14. 10.3390/socsci6010014

Hanauer, D. I., Graham, M. J., & Hatfull, G. F. (2016). A measure of college student persistence in the sciences (PITS). CBE Life Sciences Education, 15(4), ar54. 10.1187/cbe.15-09-0185

Hill, P. W., McQuillan, J., Talbert, E., Spiegel, A., Gauthier, G. R., & Diamond, J. (2017). Science Possible Selves and the Desire to be a Scientist: Mindsets, Gender Bias, and Confidence during Early Adolescence. Social sciences (Basel, Switzerland), 6(2), 55. 10.3390/socsci6020055

Hogg, Michael Al, Deborah J. Terry, and Katherine M. White. (1995). A Tale of Two Theories: A Critical Comparison of Identity Theory with Social Identity Theory. Social Psychology Quarterly, 58(4), 255–269. https://doi-org.proxy.library.emory.edu/10.2307/2787127

Huffmyer, A. S., O’Neill, T., & Lemus, J. D. (2022). Evidence for professional conceptualization in science as an important component of science identity. CBE—Life Sciences Education, 21(4), Article ar57. 10.1187/cbe.20-12-0280

Jacquez, F., Vaughn, L., Boards, A., Wells, J., & Maynard, K. (2020). Creating a culture of youth as co-researchers: The kickoff of a year-long STEM pipeline program. Journal of STEM Outreach, 3(1), 1–11. 10.15695/jstem/v3i1.02

Kim, A. Y., Sinatra, G. M., & Seyranian, V. (2018). Developing a STEM Identity Among Young Women: A Social Identity Perspective. Review of Educational Research, 88(4), 589–625. 10.3102/0034654318779957

Lucas, K. L. & Spina, A. D. (2022). The development of science identity and its implications for STEM retention and career aspirations through a research-based first year biology seminar. Journal of College Science Teaching, 52(1).

Moradi, B., Brewster, M. E., Grzanka, P. R., & Miller, M. J. (2023). The hidden curriculum of academic writing: Toward demystifying manuscript preparation. Journal of Counseling Psychology, 70(2), 119–132. 10.1037/cou0000650

Murphy, K. M., & Kelp, N. C. (2023). Undergraduate STEM students’ science communication skills, science identity, and science self-efficacy influence their motivations and behaviors in STEM community engagement. Journal of microbiology & biology education, 24(*1*), e00182–22. 10.1128/jmbe.00182-22

NGSS Lead States. (2013). Next Generation Science Standards: For states, by states. National Academies Press.

Otero, C. E., Osinski, V., & Mattison, K. A. (2022). The untapped potential of early career researchers in academic publishing: Lessons learned from the Journal of. Learned Publishing, 35(*3*), 393–399. 10.1002/leap.1470

Robinson, K. A., Perez, T., Carmel, J. H., & Linnenbrink-Garcia, L. (2019). Science identity development trajectories in a gateway college chemistry course: Predictors and relations to achievement and STEM pursuit. Contemporary Educational Psychology, 56, 180–192. doi: 10.1016/j.cedpsych.2019.01.004

A. K., Roseth, C. J., & Linnenbrink-Garcia, L. (2018). From science student to scientist: Predictors and outcomes of heterogeneous science identity trajectories in college. Developmental Psychology, 54(10), 1977–1992. doi: 10.1037/dev0000567Robnett, R. D., Chemers, M. M., & Zurbriggen, E. L. (2015). Longitudinal associations among undergraduates’ research experience, self-efficacy, and identity. Journal of Research in Science Teaching, 52(6), 847-867. 10.1002/tea.21221

Ross, M. B., Glennon, B. M., Murciano-Goroff, R., Berkes, E., Weinberg, B. A., & Lane, J. I. (2022). Women are credited less in science than men. Nature, 608, 135–145. 10.1038/s41586-022-04966-w

Smith, B. (2013). Mentoring at-risk students through the hidden curriculum of higher education. Lexington Books.

Vries, M., Land-Zandstra, A., & Smeets, I. (2019). Citizen scientists’ preferences for communication of scientific output: A literature review. Citizen Science: Theory and Practice, 4(1), 1–13. 10.5334/cstp.136

Wang MT, Degol J. (2013) Motivational Pathways to STEM Career Choices: Using Expectancy-Value Perspective to Understand Individual and Gender Differences in STEM Fields. Dev Rev. Dec 1;33(4):10.1016/j.dr.2013.08.001. doi: 10.1016/j.dr.2013.08.001. PMID: 24298199; PMCID: PMC3843492.

Watts-Taffe, S., Vaughn, L. M., & Jacquez, F. (2022). Disciplinary literacy in action: High school students doing research for change. Journal of Adolescent & Adult Literacy, 65(4), 331–341. 10.1002/jaal.1211

