## Supplemental 1 for "Writing for identity? Exploring the motivations of pre-college students to participate in science publication"

**Supplemental Material: [JOURNAL} Author Pre-Survey Questions**

**Consent** Q16: Emory University Online Consent for a Research Study [Full consent form text provided to participants]

**Demographic Information**

Q1: How did you first find out about [JOURNAL}?

- Response options: Teacher, Friend, Google, Science fair, Other

Q17: What race do you primarily identify as?

- American Indian or Alaska Native
- Asian
- Black or African American
- Latinx
- Middle Eastern
- Native Hawaiian or Other Pacific Islander
- White
- Prefer not to answer
- Other, please specify

Q18: What gender do you most identify with?

- Male
- Female
- Non-binary
- Prefer to self-describe
- Prefer not to answer

Q2: What grade are you currently in? [Open response]

**Research Context**

Q23: For the period in which you performed the research that you wrote about, what type of school did you attend?

- Public school
- Private school
- Charter school
- Home school
- College
- Other, please specify below

Q4: In what setting did you perform the majority of the research that you wrote about?

- At home
- In the school that I attend
- University/college laboratory
- Other, please specify below

Q5: Who helped you perform the research that you wrote about? Select all that apply.

- Science teacher
- Non-science teacher
- College/university professor
- Parent
- Friend or classmate
- Mostly performed independently
- Other, please specify below

Q7: Who helped you write the paper that you submitted? Select all that apply.

- Science teacher
- Non-science teacher
- College/university professor
- Parent
- Friend or classmate
- Mostly performed independently
- Other, please specify below

Q19: What is the highest degree earned by one of your parents?

- Less than high school equivalent
- High school diploma or GED
- Some college
- Associate's degree
- Bachelor's degree
- Some graduate school
- Master's degree
- Professional degree (PhD, JD, MD, LLM, etc)
- Unsure

Q8: Do you have a family member who is a scientist or engineer and helped you with the research?

- Yes
- No

Q31: Did you attend a [JOURNAL} mini-Ph.D. program?

- Yes
- No

**Writing Experience and Beliefs**

Q9: The following statements are about your experiences writing the paper you submitted to [JOURNAL}. The paper you submitted to [JOURNAL} is considered a primary science research paper (or primary paper) because you are describing the experiments that you performed. Rate the degree to which you agree or disagree with each statement:

Q9_1: Before deciding to write a paper for [JOURNAL} I was familiar with the process of writing a primary science research paper

- Scale: 1 (Strongly disagree) to 5 (Strongly agree)

Q9_2: The [JOURNAL} online guides were helpful during my writing process

- Scale: 1 (Strongly disagree) to 5 (Strongly agree)

Q9_3: The writing process was challenging for me

- Scale: 1 (Strongly disagree) to 5 (Strongly agree)

Q9_4: Submitting a paper through [JOURNAL} helped me think more carefully about the scientific process

- Scale: 1 (Strongly disagree) to 5 (Strongly agree)

Q9_5: Writing about my research helped me understand the science better

- Scale: 1 (Strongly disagree) to 5 (Strongly agree)

Q9_6: I am confident as a scientific writer

- Scale: 1 (Strongly disagree) to 5 (Strongly agree)

Q13: If you had experience with the primary literature prior to [JOURNAL} please describe that experience [Open response]

**Research Beliefs and Self-Efficacy**

Q14: The following statements are about your views in doing scientific research, such as the research you wrote your paper about. Considering the research that you wrote about, rate the degree to which you agree or disagree with each statement. [Scale: 1 (Strongly disagree) to 5 (Strongly agree) for all items]

Q14_1: My research will help to solve a problem in the world

Q14_2: My findings are important to the scientific community

Q14_3: The research question I worked on was important to me

Q14_4: I faced challenges in completing my research project

Q14_5: I was able to overcome challenges posed by my research project

Q14_6: The findings of my research project and paper gave me a sense of personal achievement

Q14_7: I am confident that I can generate a research question to answer

Q14_8: I am confident that I can figure out what data/observations to collect for a research project

Q14_9: I am confident that I can create explanations for the results of my science

Q14_10: I am confident that I can use scientific literature and reports to guide my research

Q14_11: I have a strong sense of belonging to the community of scientists

Q14_12: I have come to think of myself as a scientist

Q14_13: I feel like I belong in the field of science

**Peer Review and Publication Beliefs**

Q11: The following statements are about the general peer review process that is part of publishing a paper in an academic journal like [JOURNAL}. The paper you submitted to [JOURNAL} will undergo peer review by 3-4 senior scientists with expertise in the scientific topic of your paper. Rate the degree to which you agree or disagree with each statement. [Scale: 1 (Strongly disagree) to 5 (Strongly agree) for all items]

Q11_1: Peer review improves the accuracy of the science in a paper

Q11_2: Peer review improves the communication of the science presented

Q11_3: Peer review can give scientists different ways to understand their science

Q12: The following statements are about the role of publication within science. Rate the degree to which you agree or disagree with each statement. [Scale: 1 (Strongly disagree) to 5 (Strongly agree) for all items]

Q12_1: Publication is important because it helps a scientist share his or her science with a broader audience

Q12_2: Publication is important because the science in a paper could be used by another scientist in his or her project

Q30: What do you think is the relationship between writing and science? [Open response]

**Scientific Values**

Q15: Please read each description and think about how much each person is or is not like you. Check the answer that best reflects how much the person in the description is like you. [Scale: Not like me at all, A little like me, Somewhat like me, Like me, Very much like me]

Q15_1: A person who thinks discussing new theories and ideas between scientists is important

Q15_2: A person who thinks it is valuable to conduct research that builds the world's scientific knowledge

Q15_3: A person who thinks that scientific research can solve many of today's world challenges

Q15_4: A person who feels discovering something new in the sciences is thrilling

**Research Development**

Q21: How did you come up with your research idea? Select all that apply.

- I learned about the subject in class
- A teacher proposed the question
- A previous project that I performed sparked the idea
- Something I read sparked the idea
- A parent suggested the idea
- Talking with a friend or classmate
- Other, please explain

Q25: Please describe the process by which you thought of and developed your research project: [Open response]

**Motivation for Publication**

Q27: What motivated you to submit your paper to [JOURNAL}? Select all that apply.

- I want feedback on my research project
- I want to connect with other scientists
- I want to share my research with others
- I think a publication will give me a competitive advantage (eg: make my college application stand out)
- I want to learn how to write a scientific paper
- I want to learn about the peer-review process
- I want to improve my scientific skills
- I think others can benefit from my research
- I want to be recognized as a scientist
- Other, please explain

Q26: Please provide more detail on how the factor(s) you selected above motivated you to publish in [JOURNAL}: [Open response]

**Note:** Questions Q14_1 through Q14_13 were adapted from the Persistence in the Sciences (PITS) survey (Hanauer, Graham, & Hatfull, 2016). Questions about motivation (Q27) were developed by the research team. The survey was designed in 2020, and a focus group of student authors was convened to receive feedback on their interpretation of the questions.
